# Gene prioritization based on random walks with restarts and absorbing states, to define gene sets regulating drug pharmacodynamics from single-cell analyses

**DOI:** 10.1101/2021.02.19.431974

**Authors:** Augusto Sales-de-Queiroz, Guilherme Sales Santa Cruz, Alain Jean-Marie, Dorian Mazauric, Jérémie Roux, Frédéric Cazals

## Abstract

Prioritizing genes for their role in drug sensitivity, is an important step in understanding drugs mechanisms of action and discovering new molecular targets for co-treatment. To formalize this problem, we consider two sets of genes *X* and *P* respectively composing the predictive gene signature of sensitivity to a drug and the genes involved in its mechanism of action, as well as a protein interaction network (PPIN) containing the products of *X* and *P* as nodes. We introduce `Genetrank`, a method to prioritize the genes in *X* for their likelihood to regulate the genes in *P*.

`Genetrank` uses asymmetric random walks with restarts, absorbing states, and a suitable renormalization scheme. Using novel so-called saturation indices, we show that the conjunction of absorbing states and renormalization yields an exploration of the PPIN which is much more progressive than that afforded by random walks with restarts only. Using MINT as underlying network, we apply `Genetrank` to a predictive gene signature of cancer cells sensitivity to tumor-necrosis-factor-related apoptosis-inducing ligand (TRAIL), performed in single-cells. Our ranking provides biological insights on drug sensitivity and a gene set considerably enriched in genes regulating TRAIL pharmacodynamics when compared to the most significant differentially expressed genes obtained from a statistical analysis framework alone. We also introduce *gene expression radars*, a visualization tool to assess all pairwise interactions at a glance.

`Genetrank` is made available in the Structural Bioinformatics Library (https://sbl.inria.fr/doc/Genetrank-user-manual.html). It should prove useful for mining gene sets in conjunction with a signaling pathway, whenever other approaches yield relatively large sets of genes.

## 1 Introduction

### 1.1 Single-cell differential expression analyses

Gene differential expression analyses quantify the changes in gene expression levels between tested experimental conditions. Although gene functions can often be derived from this type of analyses, the associations can be confounded with incidental gene induction. Therefore interpreting differential expression data with gene set enrichment analysis (GSEA) and pathway analysis can be misleading without attentive curation. Single-cell differential expression analyses have elevated the issue, where gene differential expression between experimental groups is hindered by gene expression variability between cells.

Although cell-to-cell variability in gene expression is typically overlooked in most analyses, we and others have observed that these differences between clonal cells, can impact the overall cell population response to a stimulation [37, 32, 26]. Originating from stochastic processes such as transcription initiation, cell-to-cell variability gives rise to an equilibrium of co-existing cellular states within an isogenic cell population [12, 33]. The different phases of the cell cycles are one illustration of cell states that robustly proportioned resting cell populations [16], but other functional cell states can be phenomenologically evidenced by the fractional response to cytotoxic cancer drugs for example (IC50, Emax>0, [35, 27]). However, most single-cell technologies are still unable to access meaningful cell information within clonal populations once cell cycle states signatures are regressed out, and important functional cell states remain confounded in gene expression noise [26]. Apart from cell cycle genes, one hypothesis for the undetected (or unmeasurable) differences in seemingly homogeneous cell populations is that they contain cells in a wide variety of possible cell states that predisposed them to a number of responses or functions such as cell death, impairing pathway enrichment analyses. To recover the molecular determinants of clonal cells response to cancer drugs from the measured gene expression variability, we recently designed a same-cell functional pharmacogenomics approach, named `fate-seq`, that couples prior knowledge on the cell state (predicted drug response) to the transcriptomic profile of the same cell [26]. With our same-cell approach, we could reveal the molecular factors regulating the efficacy of a drug treatment, from differential expression analyses of one sample of isogenic cells with no gene induction. Although genes differential expressions can now be linked to their functional role in drug response using `fate-seq`, prioritizing genes as best potential targets for co-treatment remains a difficult task.

### 1.2 Diffusion distances

Indeed, gene prioritization and protein function prediction are challenging due to the small-world nature of interaction networks [45, 1], in particular. To go beyond analysis using the direct neighbors of a node or shortest paths, the value of diffusion distances has been recognized long ago.

Based on the correlation between the expression profile and the phenotype, as well as (diffusion) distances, various similarity measures between genes have been studied [23]. The *Diffusion State Distance* (DSD) was defined as the L1 norm of *m* walks (RW) of *k* steps [5]. The DSD was shown to be more effective than shortest-path distance to transfer functional annotations across nodes in protein-protein interaction networks (PPIN). The DSD was further extended to exploit annotations (weights) on edges, and to exploit an augmented graph incorporating specific interactions [4]. To bias the random walk towards certain nodes, a random walk restarting (RWR) at those nodes can be applied, as initially used in the context of internet surfing [29]. The stationary distribution of the strategy, called the *page rank*, depends on the restart probability vector [2]. In [43], the minimum of the page rank probability between two nodes is used to qualify the mutual affinity of two proteins in a network. RWR were also used to predict drug-target interactions on heterogeneous networks [9], and also for layered/multiplex graphs, so as to combine complementary pieces of information [41].

A related topic is the problem of ranking differentially expressed genes across two conditions [20]. Following the seminal work on gene set enrichment methods [39], the template of gene set enrichment analysis (GSEA) consist of three steps, namely computing an enrichment score for each gene, estimating its statistical significance, and performing a correction for multiple hypothesis testing. Setting aside the issue of correlations between genes, such methods combine feature selection and clustering (one cluster per cell state/condition) [15], but do not address the question of *connecting* two sets of genes using an interaction network.

Finally, yet another related challenge is pathway enrichment analysis [34]. Given a set of experimentally determined genes and a database of pathways, the goal here is to find pathways whose genes are over-represented in the gene set of interest. When pathways are known exhaustively, such analysis are sufficient to screen gene sets. If this assumption does not hold, finding *intermediates* between the gene set and the molecules in a pathway becomes mandatory.

Adding to other prioritization methods [13], `Genetrank` utilizes prior knowledge from functional cell states (transcriptomic profile of a predicted drug response), protein-protein interactions, and the expected target signaling pathway of a drug of interest.

### 1.3 Contributions

We focus on gene prioritization related to a complex phenotype, based on expression profiles from single-cell RNA-seq. Formally, let *X* be a set of proteins associated with differentially expressed genes, and *P* be a set of proteins involved in a signaling pathway. Our goal is to prioritize genes in *X* given the knowledge of genes in *P*, using an underlying PPIN, to find out those genes having a higher likelihood to regulate the pathway. Previous work on diffusion distances (RW or RWR) has three limitations in this context. First, in using hit vectors or stationary distributions, all nodes of the network contribute to the comparison of two sources. Instead, we wish to focus on nodes in *P*. Second, RWR use a bias on sources, but in our case, the sets *X* and *P* may be considered on an equal footing, which commands analysis in both directions, *i.e*. from *X* to *P* and from *P* to *X*. Third, instead of using a single restart rate [41], we study a filtration (sequence of nested sets) of genes retrieved, in tandem with so-called saturation indices revealing *accessibility scales* in the PPIN.

To accommodate this rationale, we present a novel analysis technique based on random walks with restarts and absorbing states. Recall that in a Markov chain, an absorbing state is a state which is never exited. Since stationary distributions are irrelevant in this context [11], we resort to hitting probabilities for the nodes on the target set *P*. As we show, doing so yields scores for pairs of genes in *S* × *T* and *T* × *S*, from which a ranking of genes in *X* is defined. As a case study, we use a dataset of differentially expressed genes involved in the regulation of a cancer drugs pharmacodynamics [26].

## 2 Material

### Goal: formal statement

Consider two sets of genes *X* and *P* respectively composing the predictive gene signature of sensitivity to a drug and the genes involved in its mechanism of action, as well as a protein interaction network (PPIN) containing the products of *X* and *P* as nodes. We introduce `Genetrank`, a method to prioritize the genes in *X* for their likelihood to regulate the genes in *P*.

### Biological problem

The main goal of our approach is to ameliorate the ranking of gene signatures from differential expression analyses in order to better select the genes whose products are likely to impact the phenotypic response of the cells. Using `fate-seq`, we focus on cancer cells briefly treated with the TNF-related Apoptosis Inducing Ligand (TRAIL), so for the drug response to be predicted for each cell that is then profiled by single-cell RNA sequencing [26]. In such single-cell analyses performed with isogenic cells treated together, the stable differences in gene expression between cells from the two groups, namely predicted sensitive and predicted resistant, are small and otherwise confounded in gene expression noise. In addition, the short treatment necessary for the cell response prediction does not induce a detectable genomic response (Mendeley data Dataset https://doi.org/10.17632/m289yp5skd.1 [30, 26]) that would confound the functional role of the differentially expressed genes between the two groups with gene induction, as it is often the case in other studies.

### Sets *X* and *P*

The list of differentially expressed genes obtained in this study constitutes our set *X* [26, Table S2]. The criteria used to define the two groups of single-cells compared in the differential expression analyses giving *X*, is the activation rate of caspase-8. (Caspase-8 is a protein of the receptor-mediated apoptosis pathway and of the TRAIL mechanism of action [35].) Therefore, in our case study the set *P* is a set of regulatory proteins of the extrinsic apoptosis signaling pathway.

The sets *X* and *P* are of size 320 and 49, respectively. Altogether, these sets *X* and *P* represent a unique dataset to assess the benefit of our approach in ranking genes based on their likelihood to impact drug response.

### PPIN

A number of interaction databases coexists, each with specific features, in particular a trade-off between exhaustivity and confidence. We use MINT due in particular to the compliance with the protein naming standards [7].

The PPIN was constructed from interactions downloaded from the MINT website https://mint.bio.uniroma2.it/. Only proteins with the *species* label *Homo Sapiens* were downloaded. The interacting proteins are identified in UniProtKB format. The resulting network is called the *MINT network* in the sequel. This initial graph, containing 11,672 vertices representing proteins, and 52,839 edges representing protein - protein interactions, is edited as follows. First, the PPIN being disconnected, we focus on its largest connected component, encompassing 11,427 vertices. Second, we remove all self-interactions. Third, multiple interactions between the same two proteins are collapsed into a single edge. Summarizing, we obtain a graph with 11,427 vertices and 36,526 edges.

### Sets *X* and P within the PPIN

Selected genes in the sets *X* and *P* being absent from the largest connected component of the PPIN were removed, finally obtaining sets *X* and *P* of size 227 and 41 (from sets of size 320 and 49 initially).

### Variations on the set *X*

In order to evaluate the impact of the size difference of *X* and *P* on the scores, we analyse the symmetry *H*(*I*) of Eq. (7) for instances involving sets *X*′ of varying size. Practically, for each *s* ∈ {|*P*|, 110, 220} we pick *N_r_*(= 1000) random subsets of *X*. We then compare the distributions of *H*(*I*) obtained.

## 3 Methods

We consider a set *X* of experimentally determined genes, and a set *P* of genes belonging to a pathway. We introduce methods using random walks on graphs (see below), with two sets of vertices as input, referred to as sources (*S*) and targets (*T*). Practically, we use these methods in two direction *X* ⇝ *P* and *P* ⇝ *X*.

### 3.1 Graphs

To connect *X* and *P*, we consider a PPIN whose vertices are the individual molecules, and whose edges represent pairwise interactions. Such a network is modeled by a vertex-weighted edge-weighted directed graph *G* = (*V,E*). The weight of any vertex *u* ∈ *V* is denoted *w_u_* and the weight of any edge *uv* ∈ *E* is denoted *w_uv_*. We assume that *w_u_, w_uv_* ∈ (0,1] for every *u* ∈ *V* and every *uv* ∈ *E*. In the unweighted case, we set *w_u_* = 1 and *w_uv_* = 1 for every *u* ∈ *V* and every *uv* ∈ *E*. In the undirected case, we have *uv* ∈ *E* if and only if *vu* ∈ *E*.

Let *n* = |*V*| be the number of vertices and let *V* = {*v*_1_,…,*v_n_*}. The set of out-neighbors (resp. in-neighbors) of a vertex *u* ∈ *V* is denoted 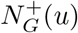 (resp. 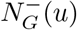).

To analyze paths between vertices of *X* and vertices of *P*, we formalize Markov chains and random walks.

### 3.2 Random walks and Markov chains with absorbing states and restart

#### Model

In the sequel, we consider two sets of vertices *S* and *T* from the graph *G*, with *S* ⋂ *T* = Ø.

We define a Markov chain for which the set of states is exactly the set of vertices *V*. The transition matrix *M* is defined as follows for every pair (*u,v*) ∈ *V* × *V*:

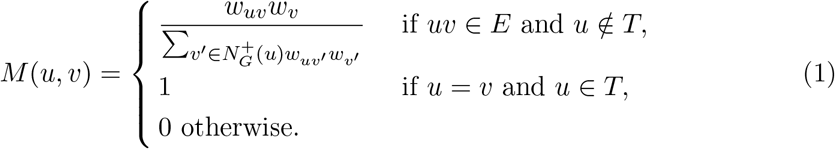

Particular cases of this construction are as follows:

- (Symmetric unweighted case) 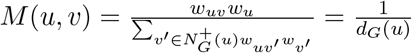 (first line of Eq. (1)).
- (Symmetric edge-weighted case) Eq. (1) also generalizes the formulae used in [4].

Recall that a state is absorbing if once reached by a walk, it is never exited. Observe that the set of states *T* is absorbing in the Markov chain defined by transition matrix *M*. We now define the Markov chain with restart from *M* and from a subset *S*′ ⊆ *S*. Intuitively, for each vertex *u* ∈ *V* \ *T* which is not a target, we add a transition to every vertex of *S*′ Formally, given a real number *r* ∈ [0,1), the transition matrix 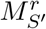 is defined as follows for every pair (*u,v*) ∈ *V*^2^.

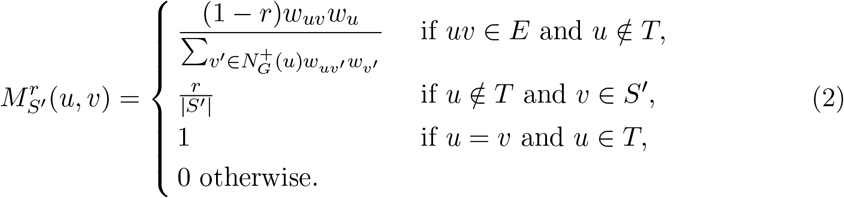

Note that restart transitions may have the same origin/destination as an existing transition in *M*. (Equivalently, in graph theoretical terms, one has two arcs between the same two vertices.) In that case, the probabilities specified in (2) should be added. The transition matrix *M* is a particular case of 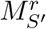 when *r* = 0. Note also that when |*S*′| = 1, one restarts to a single vertex.

##### Definition. 1 (State distribution)

*Consider an initial distribution uniform in the set S′. Given any r* ∈ [0,1) *and any S′ ⊆ S, the state distribution at each step i* ≥ 0 *is denoted*

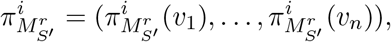

*with 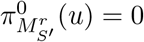 (u)* = 0 *for every u* ∉ *s′ and 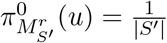 for every u* ∈ *S′*.

Under the assumption that, from every *u* ∈ *S*′ there exists a path in *G* to some *v* ∈ *T*, the limit of this vector exists when *i* → ∞, for every value of *r* ∈ [0,1). We denote it with 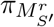. The probabilities in this vector are commonly known as hitting probabilities of the target set *T*.

Using the target set, we define:

##### Definition. 2 (hit probability vector)

*Let T* = {*t*_1_,…,*t_k_*} *be the target set. Given any r* ∈ [0,1) *and any S*′ ⊆ *S, the hit vector* 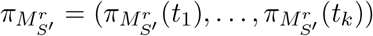 *is composed of the hitting probabilities for states of T*.

In the following, we abuse the notation writing 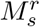. instead of 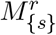. Furthermore, we will write *M^r^* instead of 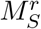. Finally, we renormalize the vectors with the hit vector of the chain without restart (*i.e*. *r* = 0):

##### Definition. 3 (hit score)

*Given r* ∈ [0,1], *define the score from s to t as*

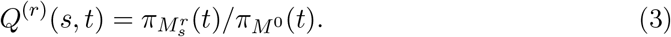

*The* hit score vector *associated with each source s* ∈ *S is (Q^(r)^*(*s,t*_1_),…,*Q^(r)^(s,t_k_)). The log score* log *Q^(r)^(s,t) is the natural logarithm of Q^(r)^(s,t)*.

Finally the rank of the score of a pair (*s, t*) ∈ *S* × *T* is defined as follows:

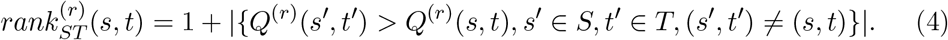

**Remark 1** *The hit score from Eq. 3 incorporates three ingredients, namely (i) a random walk with restart, (ii) absorbing states, and (iii) renormalization by the value obtained without restart. We may define other scores by removing any of these ingredients, e.g. the mechanism of absorbing states*.

#### Comparison to previous work

In this work, we aim at computing the relative importance of the targets of *T* compared to the sources of *S*. Two structural properties of interaction networks actually underlie our modifications based on absorbing states and the renormalization.

The first difficulty owes to a notion of *subsidiarity* amongst targets, meaning that if some vertices of *T* are neighbors (or very close from one another) in the graph, targets *upstream* will artificially modify the weights of targets *downstream*. Indeed, if a target is just *after* another (important) target, then its weight will be large even in the absence of direct paths from *S* (Fig. 1, target *t*_4_). In this context, absorbing states make it possible to stop the exploration process at such *upstream/ancestor nodes*, and force direct connexions from sources to *subsidiary* targets.

**Figure 1:**
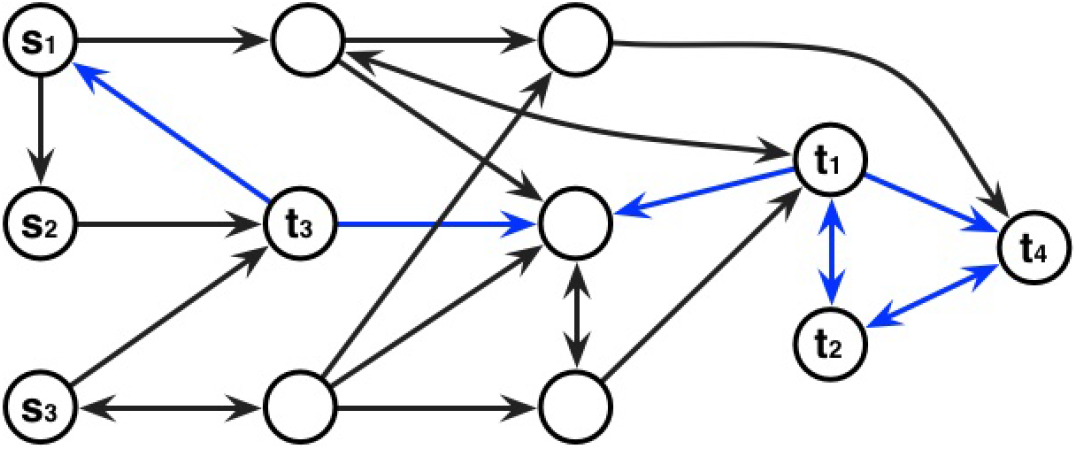
Example interaction graph and Markov chain: two structural properties motivating absorbing states and the renormalization of hit probabilities. Arcs indicate transitions in the Markov chains - transition probabilities are omitted. The set of sources is *S* = {*s*_1_,*s*_2_,*s*_3_} and the set of targets is *T* = {*t*_1_,*t*_2_,*t*_3_,*t*_4_}. When defining absorbing states, transitions corresponding to blue arcs are removed. This implies that *t*_2_ will not be highlighted because *t*_1_ or *t*_4_ must be visited before in any random walk starting from any source *s* ∈ *S*. Furthermore, the normalization aims at reducing the importance of *t*_3_ in the study (due to its high proximity to the three sources).

The second difficulty owes to the close vicinity of selected targets to sources (Fig. 1, target *t*_3_), motivating the introduction of hit scores (Def. 3). For example, if a target is a neighbor of some sources in the graph, these hit scores decrease the weights of such *direct paths*, stressing the importance of the other *non trivial* paths. One can note that this normalization can be done with other functions, e.g. a distance-based function that could be the average of the |*T*| lengths of shortest paths between every source *s* ∈ *S* and a given target *t*.

### 3.3 Scores and their symmetry

#### Scores *X* ⇝ *P* versus *P* ⇝ *X*

In order to analyze the (lack of) symmetry between paths joining *X* and *P* and vice-versa, We apply the previous construction to two settings:

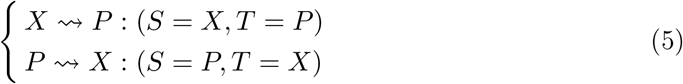

#### Instances (*PPIN, X, P*)

Using a different PPIN, a different experimental gene set *X* or a different pathway gene set *P* will affect the score *Q*^(*r*)^(*x,p*) for a given pair (*x,p*). In order to make it clearer we define an instance of execution as the triplet *I* = (*PPIN, X, P*), and we refer to the score obtained under this instance as 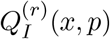. However, for conciseness, we simply denote this score *Q*^(*r*)^(*x,p*).

#### Symmetry at the gene level

Consider an experimental gene *x* ∈ *X* and a pathway gene *p* ∈ *P*. Using Eq. (3), we assess the asymmetry using the ratio of log scores

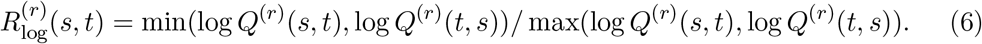

#### Symmetry at the instance level

Consider an instance *I* = (*PPIN, X, P*) ^1^. To study the symmetry at the instance level, we consider the proportion of pairs (*x,p*) ∈ *X* × *P* such that 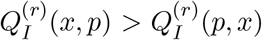. Formally, denoting **1**_*b*_ the indicator function of the boolean *b*, the symmetry of the results for *I* is given by

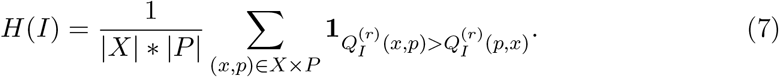

### 3.4 Genetrank, saturation indices, hits

Using the scores in the two directions *X* ⇝ *P* versus *P* ⇝ *X* (Eq. 5), we now define a ranking on the genes of *X*:

#### Definition. 4 (Average score)

*Let* 1 ≤ *τ* ≤ |*P*| *be an integer. Consider a fixed value of the restart rate r. For a source x, let the* average score *be the arithmetic mean over the top τ values* max(log *Q*^(*r*)^(*x,p*), log *Q*^(*r*)^(*p, x*)) *observed for p* ∈ *P. The gene network ranking* (`Genetrank`) *of genes in X is the ranking associated with the aforementioned average values. The set of top k genes of the ranking is denoted* 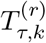.

Note that when *τ* = 1, the ranking of a gene in *X* is determined by its largest max score. Averaging scores over *τ* targets makes intuitive sense here when identifying whole connectedness to a pathway.

To assess the stability of this ranking, we proceed as follows. Consider a set of values *R* = {*r*_1_,…,*r_N_*}, sorted by increasing or decreasing value. We define the set of genes found in 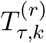 up to a given value *r_l_*, with 1 ≤ *l* ≤ *N*, by

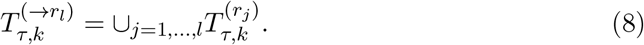

We now use this set to qualify the *speed* at which we discover the sources in *X* when increasing the upper bound on the restart rate:

#### Definition. 5 (Saturation indices for an increasing sequence of values of *r*.)

*The* saturation index *at threshold r_l_ is the fraction of sources present in 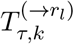 that is*:

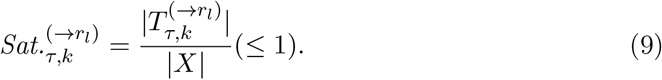

*The* relative saturation index *is the latter normalized by the value of k used*:

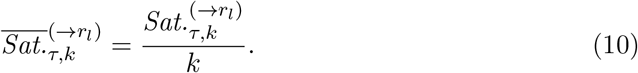

In the absence of overlap between consecutive 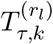, one would have 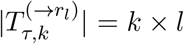. Thus, normalizing by *k* provides a measure of the overlap between consecutive sets.

We note in passing that the previous sets can be used to define how many hits in a given reference list of genes 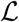 are obtained:

#### Definition. 6 (Hits)

*Consider a reference list of genes 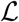. The number of hits for particular values (r, k) is the size of the set 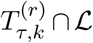. For a fixed k, we similarly consider the size of the set 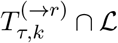*.

**Remark 2** *Saturation indices for a decreasing sequence of values of r readily generalize from Eqs. 8, 9, and 10. In fact, a larger (resp. smaller) value of r amounts to zooming in (resp. out) towards the sources*.

### 3.5 Graphical representations with radar scatter plots

We wish to rank genes from *X* using genes from *P*, exploiting the directions *X* ⇝ *P* and *P* ⇝ *X*.

#### Score radar plots

The difficulty in working with values LogCount(*x*), LogFoldChange(*x*) for *x* ∈ *X* represented in an MA plot, is that all pairs (*x,p*) get mapped onto the same point. To get around this difficulty, we associate a *radar plot* with each point *x* ∈ *X*, yielding an overall score radar scatter plot. Each *gene score radar plot* is defined as follows:

- the background of the gene radar plot is colored using a heat map indexed on the largest (*X* ⇝ *P* or *P* ⇝ *X*) log score observed for that gene. This background color makes it easy to spot the individual radar plots with high scores.
- the gene radar plot has a number of spokes equal to the top *k* (user defined) scores.
- on each spoke, two values are found, namely the scores log *Q*^(*r*)^(*x,p*) and log *Q*^(*r*)^(*p, x*).
- finally, the radar plot title is set set to the gene name accompanied by the interval of scores (log scale).

#### Score radar MA plot

Displaying all individual score radar plots in the LogCount(*x*), LogFoldChange(*x*) plane yields the so-called Score radar MA plot.

### 3.6 Implementation

We compute hit scores (Def. 3) using the C++ `Marmote` library ([19] and https://wiki.inria.fr/MARMOTE/Welcome). The whole pipeline is implemented in the `Genetrank` package of the Structural Bioinformatics Library ([6] and http://sbl.inria.fr), see https://sbl.inria.fr/doc/Genetrank-user-manual.html.

The running time of one instance *i.e*. computing the hit scores at a fixed *r*, depends on the sizes of the PPIN, and of sets *X* and *P*. These latter two sets matter most since a Markov chain is generated and then evaluated for every source. The most time-consuming work is the calculation of the hitting probabilities, which in theory can be performed exactly with matrix inversion, at a practical cost of order *n*^3^. We selected from the `Marmote` library the iterative method which approximates the result with iterations of order *m*, with *m* = |*E*| the number of edges. It is faster and also more stable in practice, since it involves only positive numbers. The stopping criterion chosen for iterations is that the *L*_1_ distance between successive iterates is less than 10^-6^.

Practically, processing one instance took a few minutes (< 5) worst-case, on a standard desktop computer (Precision 7920 Tower, 64 Go of RAM, Intel(R) Xeon(R) Silver 4214 CPU at 2.20GHz; OS: Fedora Core 32).

### 3.7 Tests: setup

#### Contenders

As already noticed (Rmk. 1), the hit score from Eq. 3 incorporates three ingredients, namely (i) a random walk with restart, (ii) absorbing states, and (iii) renormalization by the value obtained without restart. To assess the importance of the latter two ingredients, three contenders of nested complexity are considered in the sequel:

- `pr-affinity`: the minimum page rank affinity introduced in [43], using a plain random walk with restart model.
- `Genetrank-renorm`: score obtained from a random walk with restart and renormalization using Eq. 3 - but no absorbing state.
- `Genetrank-AS`: score obtained from a random walk with restart and absorbing states - but without the renormalization of Eq. 3.
- `Genetrank`: score obtained using all ingredients: random walk with restart, absorbing states, and the renormalization of Eq. 3.

#### Parameters used

The following values are used in our experiments:

- 81 values of *r*, from *r* = 0 to *r* = 0.8 by steps of 0.01,
- three values of *τ* (Def. 4): *τ* ∈ {1, 20,41}, (recall that |*P*| = 41),
- ten values of *k*: *k* ∈ {5,10,15, 20, 25, 30, 35, 40, 45, 50},
- saturation indices for *r* sorted by increasing/decreasing values (Rmk. 2).

## 4 Results

### 4.1 Genetrank and saturation indices

The saturation index makes it possible to study the variation of the size of the set of genes selected by the three contenders when *r* increases or decreases. To promote the vicinity of sources in the PPIN, we process values *r* by decreasing value (Rmk. 2). We inspect in turn the saturation, relative saturation and number of hits (Fig. 2, Fig. 3, Fig. S4, Fig. S5). We use the median value *τ* = 20 to compute average scores – see the Supplemental for the values *τ* = 1 and *τ* = 40.

**Figure 2:**
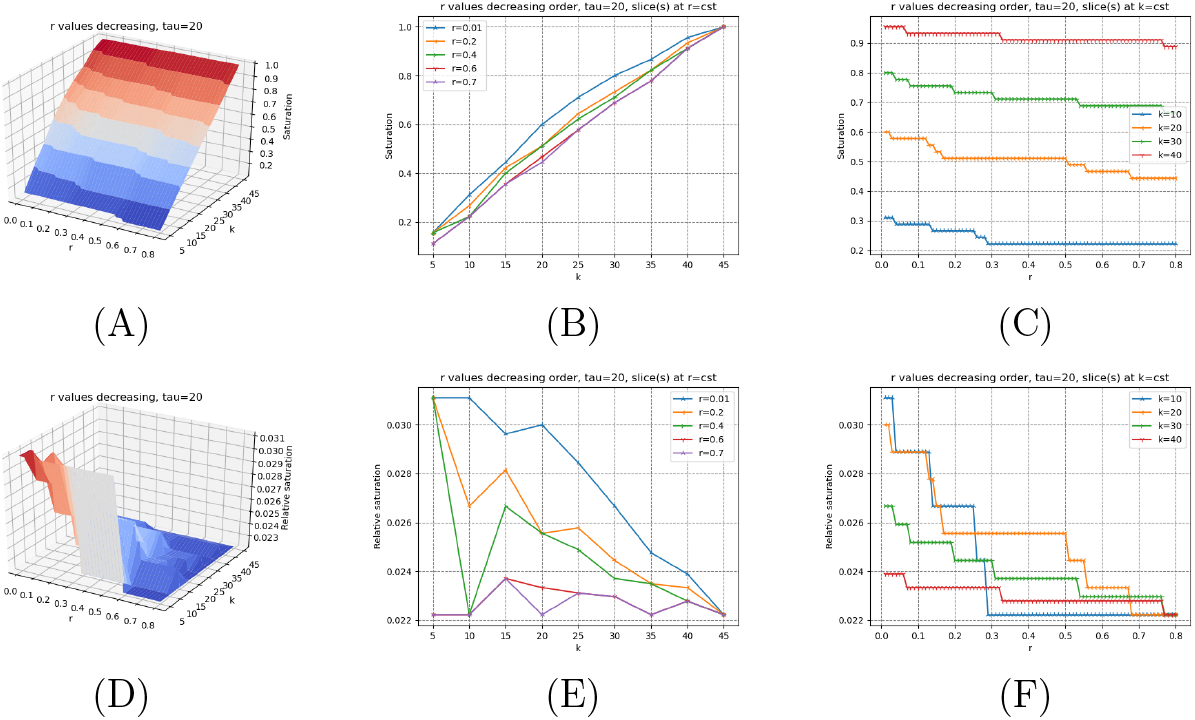
(pr-affinity) Saturation plots (Def. 5) for *τ* = 20. Values of *r* processed in decreasing order. **(A, B, C)** Saturation index and slices at *r* = *cst* and *k* = *cst* (See Eq. 9) **(D, E, F)** Relative saturation index and slices at *r* = *cst* and *k* = *cst* (See Eq. 10)

**Figure 3:**
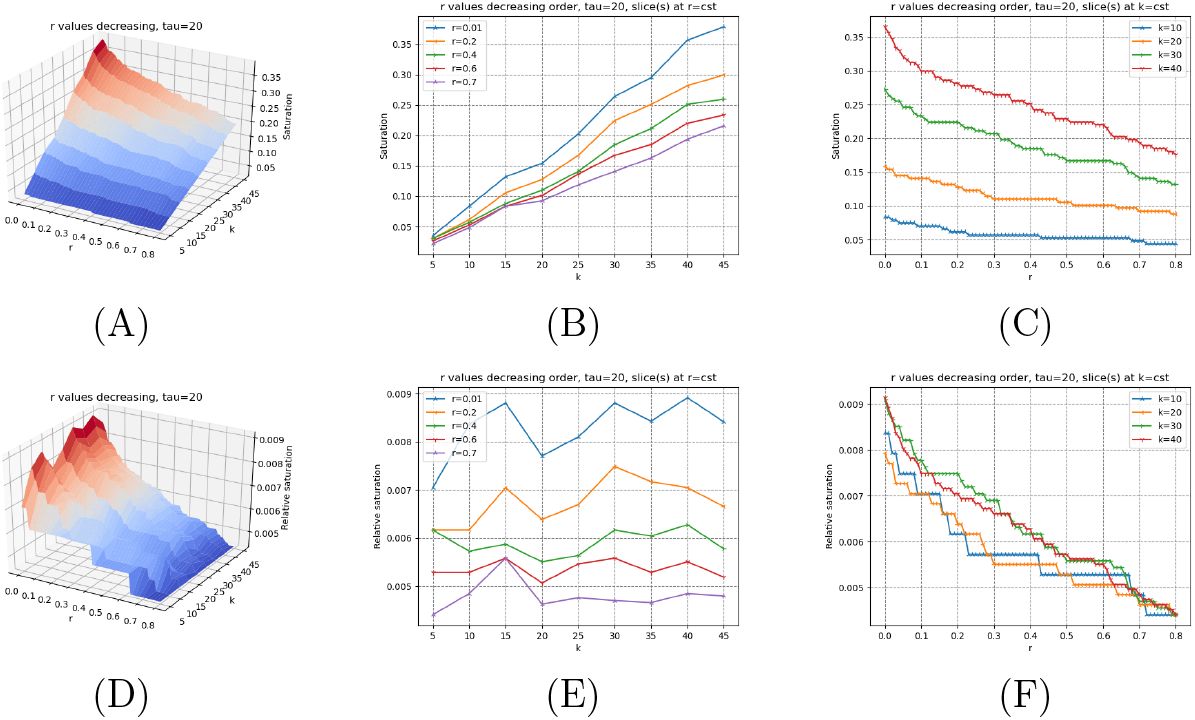
(Genetrank) Saturation plots (Def. 5) for *τ* = 20. Values of *r* processed in decreasing order. **(A, B, C)** Saturation index and slices at *r* = *cst* and *k* = *cst* (See Eq. 9) **(D, E, F)** Relative saturation index and slices at *r* = *cst* and *k* = *cst* (See Eq. 10)

The methods `pr-affinity, Genetrank-renorm`, and `Genetrank-AS` yield a very similar behavior in two respects. First, the saturation gets maximum (one) for large values of *k*, and is relatively insensitive to the value of *r* (Fig. 2(B), Fig. S4(B), Fig. S5(B)). Also, the relative saturation drops down to very small values rapidly (Fig. 2(E), Fig. S4(E), Fig. S5(E)). In sharp contrast, the maximum saturation yielded by `Genetrank` is equal to ~ 0.4, and shows a marked dependence on the restart rate (Fig. 3(B)). The relative saturation is also much more sensitive to the value of *k* used, with a coherent behavior as a function of *k* at fixed values of *r* (Fig. 3(E,F)).

These observations stress the specificity yielded by the combination of renormalization and absorbing states in `Genetrank`. To further these insights, we proceed with a more detailed analysis of the incidence of the parameters *r* and *τ*:

- (*r*) The restarting rate *r* defines the bias towards sources. Whatever the value of *k*, in processing values of *r* in decreasing values, we observe that the saturation increases when *r* approaches zero (Fig. 3(C)). The slope of the curves increases for values of *r* ≤ 0.1, showing that for such values of the restart rate, the random walks get to explore a larger region of the PPIN. In `Genetrank`, larger values of the restart rate are therefore instrumental in promoting a more specific exploration.
- (*τ*) Increasing *τ* consists of averaging on more targets. This averaging yields a marked increase of the saturation (whatever the value of *k*), especially for small values of *r* (Fig. 3, Fig. S6, Fig. S7).

Let us now consider the relative saturation (Fig. 3(D,E,F); Fig. S6(D,E,F), Fig. S7(D,E,F)). Sections of the surface at *r* = *cst* yield a monotonic behavior, that is the smaller the restart rate the larger the relative saturation. (We also note that for the first value of *r* processed, that is *r* = 0,8, the relative saturation is always equal to 1/|*X*| ~ 0.0044.) Interestingly, slices at *r* = *cst* are not monotonic. The valleys crossed on such slices show that increasing *k* increases the saturation but not necessarily the relative saturation. This may be interpreted as *accessibility scales* in the PPIN.

### 4.2 Evaluating the effect of varying restart rates on scores, using experimentally validated gene hits

Consider the list of reference genes (and their products) that had been experimentally validated following the single-cell functional genomics approach using predictive cell dynamics [26] O15304 (SIVA1), P78537 (BLOC1S1), P0CW18 (PRSS56), P53007 (SLC25A1), Q9Y2X8 (UBE2D4), DNM1L(O00429). Note that Q6UW78(UQCC3) cannot be considered here, as it is not present in our PPIN of interest. We compute the number of genes from this list found in the sets 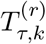 and 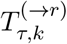.

We compare the results of the four methods (Fig. 4, Fig. 5, Fig. S8, Fig. S9). Consistent with the analysis in the previous section, the methods `pr-affinity` and `Genetrank-renorm` are rather non specific, with a number of hits yielded by 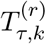 essentially constant whatever the value of *r* at a fixed value of *k*, whatever the value of *τ* (Fig. S4, Fig. S8). The method `Genetrank-AS` shows a more contrived behavior, with the same number of genes uncovered (i.e. 5) for small values of *r*, yet more variations when varying *r* at fixed values of *k* (Fig. S9).

**Figure 4:**
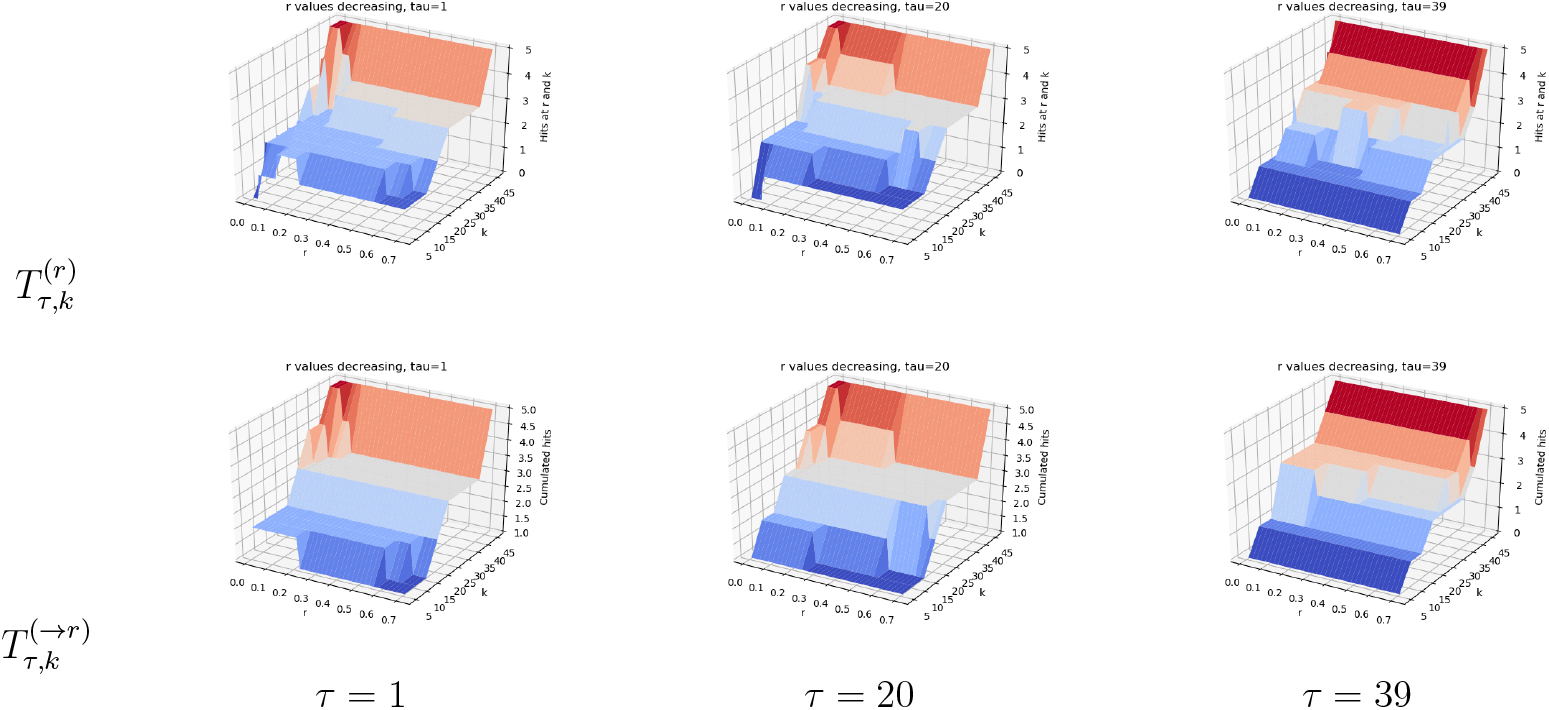
(pr-affinity) Hits (Def. 6) for the list of reference genes O15304 (SIVA1), P78537 (BLOC1S1), P0CW18 (PRSS56), P53007 (SLC25A1), Q9Y2X8 (UBE2D4), DNM1L(O00429).

**Figure 5:**
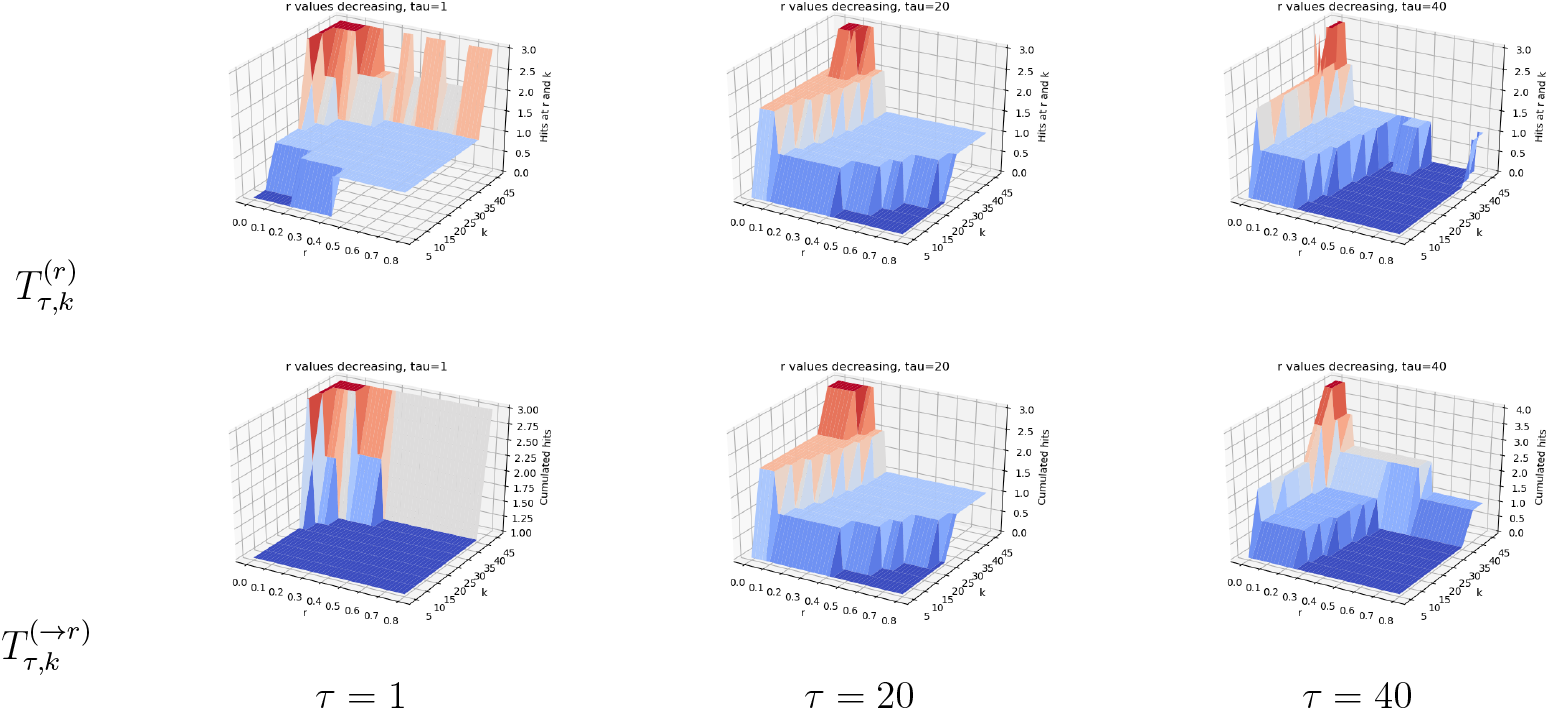
(Genetrank) Hits (Def. 6) for the list of reference genes O15304 (SIVA1), P78537 (BLOC1S1), P0CW18 (PRSS56), P53007 (SLC25A1), Q9Y2X8 (UBE2D4), DNM1L(O00429).

**Figure 6:**
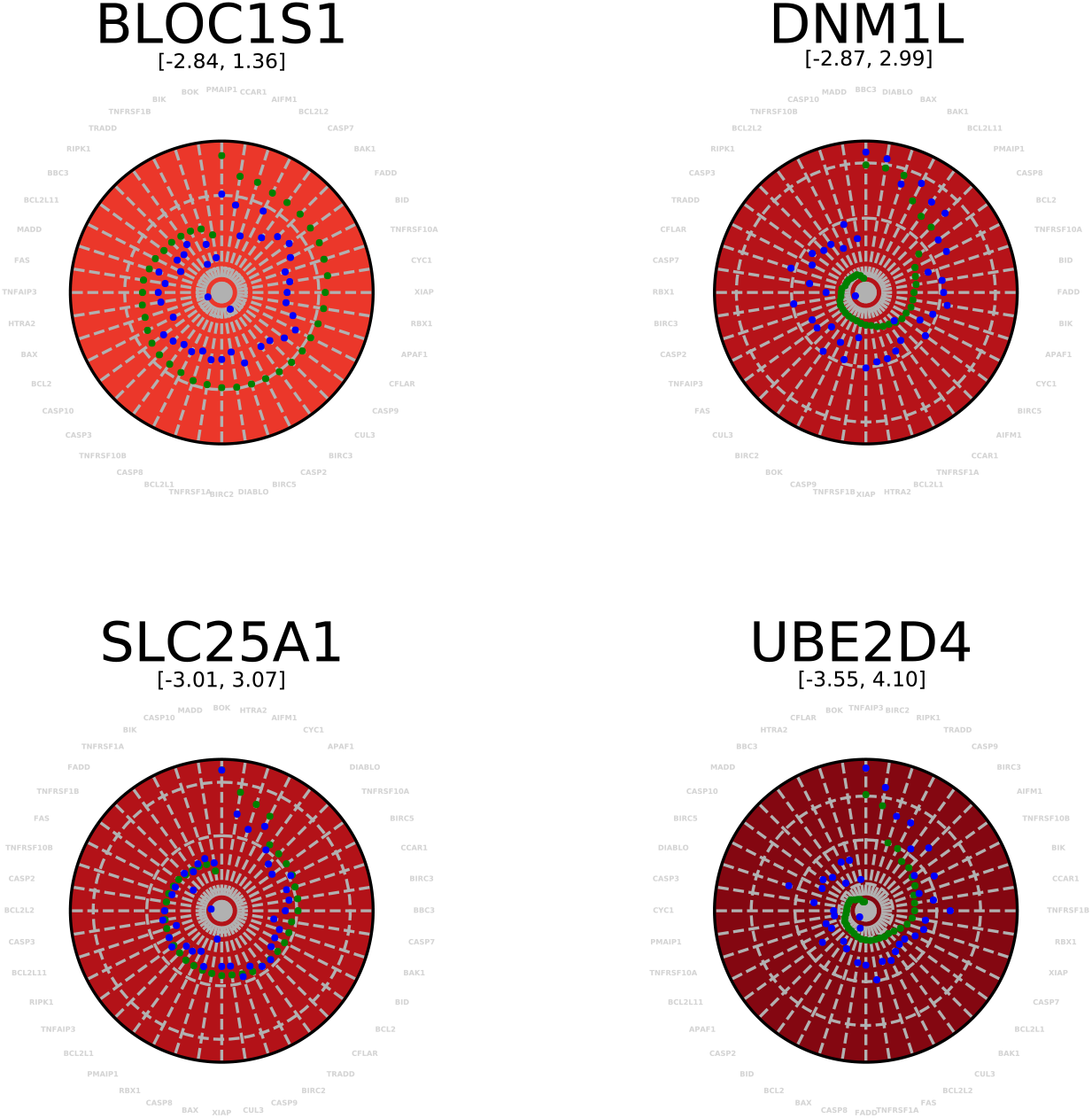
Radar plots for four experimentally validated genes. The range of values covers the range of log scores observed. The two bullets on a spoke read as follows: blue dot: direction *PX* ⇝; green dot: direction *XP* ⇝.

**Figure 7:**
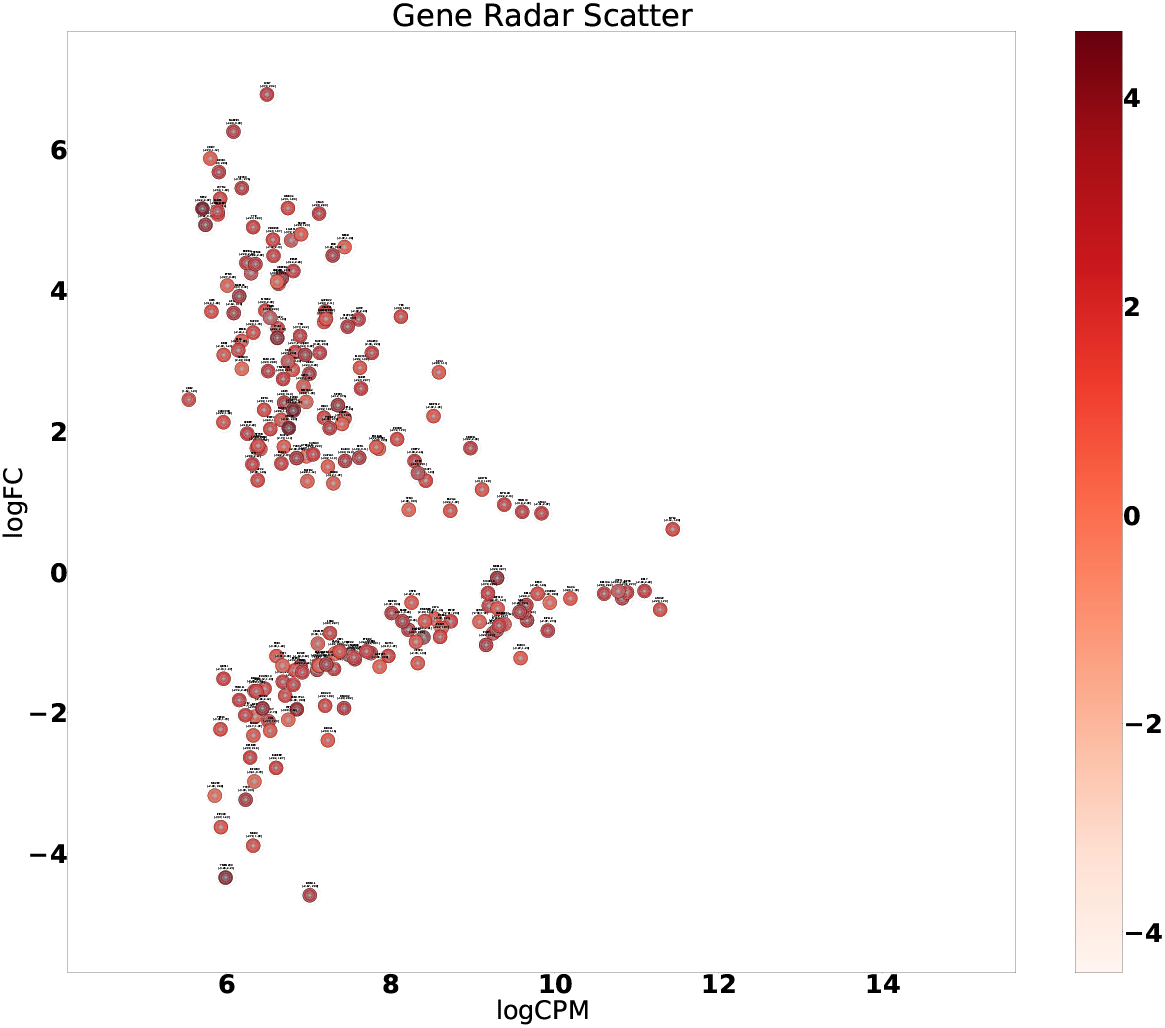
Score radar scatter plot. The individual radar plots (Fig. 6) are assembled in a MA plot.

The method `Genetrank` goes one step beyond, namely discovers genes more selectively when increasing *k* and changing the restart rate (Fig. 5). Small values of *τ* allow retrieving three genes using fewer values of *r*, clearly showing the specificity/robustness of nodes of *X* highly ranked with a large restart rate. Conversely, using larger values of *τ* in conjunction with smaller values of *r* (larger excursions in the PPIN) allows reporting four genes in total. Except for *τ* = 1, using large values of *r* requires larger values of *k* to retrieve the known genes: for *τ* = 20 and *τ* = 41, a single gene is retrieved when *r* > 0.5.

These experiments show the progressiveness yielded by `Genetrank` is exploring the PPIN, as opposed to avoid the fast *mixing* observed in `pr-affinity` and `Genetrank-renorm`, and to a lesser extent `Genetrank-AS`. Methods without absorbing states *see* the whole PPIN in a more homogeneous way, due to its *small-world* nature. This behaviour is even more pronounced in our case, as the target genes form a pathway. Indeed, a random walk reaching one such node is likely to discover its neighbors right after (See *Comparison to previous work*, Sec. 3.2.) When a absorbing states are used, the random walks halts, and the discovery of neighbors does not take place. The introduction of absorbing states therefore appears crucial to localize the exploration of connexions, in conjunction the choice of the restart rate *r* - generally taken to *r* = 0.7 in previous work - see [41] and citations therein.

### 4.3 Symmetry analysis on a per-source basis: radar plots

The symmetry ratio *H*(*I*) is a global assessment based on all pairs in the Cartesian product *X* × *P*. For an assessment of the asymmetry on a per gene basis, we resort to radar plots (Sec. 3.5). Example radar plots for genes experimentally validated are provided in the supporting information (Fig. S6).

### 4.4 Biological analysis

#### Differentially expressed genes

The set of differentially expressed (DE) genes obtained from the single-cell RNA-seq analyses chosen here, allows comparing transcriptomic profiles between single cells of an isogenic population, grouped by their predicted drug response [26]. Although this response prediction helped increase the meaningful expression signal obtained by differential analysis using the `edgeR` likelihood ratio test framework [25], the most significant differentially expressed genes (false discovery rate FDR < 0.1 and ļlog2(FC)ļ > 2) still constitute a list of more than 60 genes, defying the expected practical set of potential targets that would serve to design co-therapeutic strategies increasing overall treatment efficacy. In addition, ranking only on differential expression might underestimate a gene for its potential function as regulator of the pathway overall activity. As an example, among these gene hits, we have observed that, at equal distance (the shortest path between the source and the target gene caspase-8), the noisier the gene expression was, the larger the effect a gene perturbation had on cell death, and at comparable expression variability: the longer the shortest path, the larger the effect [26]. We reasoned that diffusion distances could also ameliorate ranking of cell-to-cell differential expression analyses.

We use `Genetrank` to sort genes for their connectedness to molecular factors regulating the signaling pathway triggered by the drug of interest (TRAIL). We intersect the list of 65 most significant differentially expressed genes obtained by `edgeR`, with the set of genes obtained with `Genetrank` (*k* = 50, range of values of *r*: 0..0.8, *τ* = 41; Eq. (8) and Fig. S7), resulting in 18 genes (Table S1). This gene shortlist presents a number of valuable advantages over the list of significant differentially expressed genes, as it becomes more manageable for experimental validation, and drug target discovery. In addition we found that this shortlist contained genes that had been previously reported to regulate cell death and importantly, it was enriched in genes that we had experimentally validated for having an effect on TRAIL response.

#### Previously validated genes

Out of the 18 genes (Table S1) we prioritized for their likelihood of having an effect on the drug mechanism of action (MoA) using `Genetrank`, we found 4 genes that we had previously validated experimentally, namely *BLOC1S1, DNM1L, UBE2D4, SLC25A1*. This result indicates that we successfully enriched the list with gene hits that have functional relevance as co-drug targets. Indeed, among the target genes shortlisted solely based on statistical criteria (differential expression analyses between predicted resistant and predicted sensitive cells to TRAIL), only *DNM1L*, had previous reports either on TRAIL sensitivity. Indeed, *DNM1L* encoding the dynamin-1-like protein DRP1, involved in mitochondrial division and apoptosis, has been reported to increase TRAIL sensitivity regardless of the co-treatment strategy (recruiting DRP1 at mitochondrial membrane, or inhibiting DRP1), which lead the authors to suggest that DRP1 might not be the target of mdivi-1, its originally reported inhibitor [21, 44]. In our experimental screen, cells that were predicted resistant to TRAIL based on low caspase-8 early activation rates, showed increased *DNM1L* expression levels [26]. Also in this recent study, we could show that *DNM1L* over expression reduced caspase-8 activation and cell death in response to TRAIL, and consistently, that DRP1 or dynamins inhibition (using mdivi-1 or dynasore), both increase caspase-8 activation and cell death. All together, these results suggest that in addition to the pro-apoptotic role of DRP1 at the mitochondrial membrane, DRP1 could play an anti-apoptotic role at receptor level on caspase-8, further validating the approach presented here and the relevance of `Genetrank`.

#### Novel genes

`Genetrank` also puts forward some genes that were initially further down the list and therefore in an unfavorable position to command gene validation or functional studies. JADE1 for example, has been shown to promote apoptosis in renal cancer cells [47], and MUL1 [31, 46], which should motivate further experimental investigations.

## 5 Discussion

### Method

This work presents a gene prioritization method, `Genetrank`, which can be coupled with single-cell functional genomics approaches to rank the drug sensitivity of a set of genes *X* for their likelihood to regulate the cell signaling pathway *P* targeted by the drug of interest. While diffusion based distances have been used for several problems in interaction network analysis, the `Genetrank` introduces several refinements. The first one is to use random walks with restarts and absorbing states to focus on certain nodes of the PPIN. The second one is to exploit the asymmetry of random walks from *X* to *P* and *P* to *X* across the PPIN. The third one is to use a whole set of restart rates to define a filtration (sequence of nested sets) of genes, using *saturation indices*. The analysis yielded by saturation indices show the ability of `Genetrank` to progressively unveil regions of the PPIN, thanks to absorbing states and our renormalization scheme. Instead, classical methods based on random walks (with or without restarts) *see* the whole network in a more homogeneous fashion due to its small-world nature. In this context, absorbing states *force* the evaluation of direct connexions between sources and targets, and the renormalization scheme makes it possible to tone down large weights due to the proximity between sources and selected targets. Altogether, these modifications allow `Genetrank` at various restart rates to progressively explore the network, and incrementally investigate interactions within a pathway. For gene prioritization, our novel ingredients make it possible to perform a delicate study of the interplay operating between the different parameters defining the RW, providing a stratification of genes of *X* according to their *proximity* to genes in *P*.

### Biology

As a case-study, we used `Genetrank` with a TRAIL-sensitivity gene signature obtained from the single-cell functional genomics approach using predictive cell dynamics called `fate-seq` [26]. Here, we show that we could enrich the most significant differentially expressed genes between predicted sensitive and resistant cells, with genes that were experimentally validated for increasing drug treatment efficacy (or previously described as doing so).

The nature and design of large transcriptomic profiling experiments (single-cell and bulk) and their analyses, impose a number of limits on the biological interpretation of gene expression analyses, especially in the context of understanding the determinants of drug sensitivity. Firstly, the differential analyses between cell types of varying drug sensitivities can be confounded with bystander genes (regarding their role in the drug MoA). Secondly, within a cell type, differential analyses between treated versus untreated samples lacks specificity over genes at the origin of the cell response versus the genes induced by the drug in resistant cells (not to mention that sensitive cells are rarely recovered in experiments). Moreover, the subsequent analyses such as gene set enrichment analysis and pathway-based analyses [39, 40, 42], dependent on prior knowledge of the gene functions and their interactions, often determined with the aforementioned experimental designs. Gene annotations themselves (from Gene Ontology (GO) database for example) might hinder gene-based drug sensitivity predictions, by introducing biases related to errors or incompleteness due to unknown function, protein moonlighting [36], or technical and biological inherent limits [24, 38]. Yet, gene expression remain a central piece of data in drug sensitivity prediction [10, 14], providing successful use with cancer cell lines to discover gene involved in drug resistance [3] with computational methods using prior knowledge [22, 28, 8, 17, 18]. Although some analyses performed these tasks on basal gene expression [13], which aim at harnessing cell states at the origin of drug response (as opposed to drug-perturbed gene expression studies), all pursue cell lines comparative profiling.

However, profiling cell lines remains ineffective with respect to determining the molecular factors involved in the incomplete -or fractional-response of one cell line, due to intrinsic drug resistance of non-genetic origins (a phenomenon observed for all drugs at their IC50 in all cell lines). Both single-cell genomics and single-cell drug response analyses [35] have revealed a range of heterogeneity within isogenic cell populations (within one “cell type”) that has not been fully comprehended up to now, for technical reasons [26]. And the natural gene expression variability of isogenic cells may be referred to as gene expression noise only because of the actual deficiency in specific gene sets that define cell states such as drug-sensitive or-resistant state (as opposed to a drug-induced state in the resistant cells), that could inform on genes likely to perturb the MoA of the drug of interest. Therefore, single-cell experimental methods to determine the MoA-perturbing genes remain critical to increase treatment efficacy (or reduce treatment toxicity).

Our approach utilizes predicted drug response knowledge from `fate-seq` to rank genes among MoA-perturbing gene signature and associate prior knowledge from proteinprotein interaction networks to favor protein that are connected to the targeted signaling pathway, which may also reveal novel biological activities.

### Future work

Our results suggest that combining same-cell functional pharmacogenomics screens such as `fate-seq`, with gene prioritization technique described here, are promising novel methods to improve gene definitions in GO with respect to their association to novel drug efficacy gene signatures, and help revealing the most effective co-targets for combination therapy.

From the theoretical standpoint, our gene prioritization strategy poses a fundamental question. Given a pair of genes highlighted (one source in *X*, one target in *P*), an open question is to find sets of intermediates nodes accounting for the hit scores used. Indeed, such intermediates could be used to delineate the biochemistry of interactions (enzymes, non covalent interactions, etc), paving the way to more quantitative models involving reaction rates and/or affinity constants.

## Acknowledgments

We acknowledge the support of (1) the Investissements d’Avenir UCA JEDI project, ANR-15-IDEX-01; (2) the 3IA Côte d’Azur Investments in the Future project managed by the National Research Agency, ANR-19-P3IA-0002; (3) the INCa Plan Cancer Biologie Des Systèmes, ITMO Cancer (proposal IMoDRez, no.18CB001-00).

## 7 Supporting information: results

### 7.1 Individual scores and their symmetry

#### Global analysis

We first notice that the scores *Q*^(*r*)^(*x,p*) and *Q*^(*r*)^(*p, x*) for all pairs *X* × *P* tend to be sharply peaked near the origin, especially for small values of the restart rate *r* (Section 7.1, Fig. S1).

To study the symmetry of scores, we resort to scatter plots whose *x* and *y* axis are the ranks of scores (Eq. 4), and values displayed are the scores log(*Q*^(*r*)^(*x,p*)), log(*Q*^(*r*)^(*x,p*)) and their difference log(|*Q*^(*r*)^(*x,p*) — *Q*^(*r*)^(*p,x*)|) (Fig. S2).

Upon inspecting these plots by varying *r*, the following appears:

- (Scatter plots, scores for *X* ⇝ *P* i.e. first column) Plotting the value for ST results in a narrow vertical band of large values - consistent with the fact that the histogram of log values is sharply peaked near zero yielding large negative logs.
- (Scatter plots, scores for in *P* ⇝ *X* i.e. second column) Likewise, plotting the value for *X* ⇝ *P* results in a narrow horizontal band, but we notice a higher density for large ranks in TS.
- (Comparing rows) Increasing the restart rate widens the range of scores (Fig. S1), which in turn stresses the aforementioned vertical and horizontal bands.
- The previous observations are combined on the difference plot (Fig. S2 (Right column)), and are especially salient for *r* = 0.3. It indeed appears that large values of the log of the difference are only obtained for a small rank in *X* ⇝ *P* (large score in *X* ⇝ *P*) or in *P* ⇝ *X* (large score in *P* ⇝ *X*); moreover, large negative values of the log are not observed near the origin, which shows that large scores in *X* ⇝ *P* and *P* ⇝ *X* are not observed concomitantly except for ranks ≤ 50.

This lack of symmetry of scores is a strong indication that paths joining *x* to *p* have significantly different features from those joining *p* to *x*, in particular in terms of high degree vertices. Every path from *x* to *p* is also a path from *p* to *x*. However, a high degree vertex which appears early in the path from *x* to *p* appears late in the reverse path. Such high degree vertices yield many alternative paths, which are more competitive with each other in the *x* to *p* direction.

#### Incidence of the size of *X* on scores and their symmetry

We study the incidence of the sizes |*X*|, |*P*| on the symmetry score ratio *H*(*I*) of Eq. (7). To do so, we pick random subsets *X*′ ⊂ *X* of size {|*P*|, 110,220}, performing 1000 repeats for each value (Fig. S3).

First, considering the statistics for the symmetry ratio *H*(*I*), despite seemingly related values for the mean and std deviation of the statistic *H*(*I*) for the two restart rates *r* = 0.01 and *r* = 0.3, the p-value of the non-parametric two-sample test used shows a strong evidence to reject the equality of distributions. Second, *H*(*I*) displays a marked dependence on the size of *X*, which is related to two facts. On the one hand, increasing the size of *X* does not affect the score in the direction *X* ⇝ *P*; however, the hitting probabilities in the direction PX are getting *diluted*. On the other hand, due to the small-world nature of the graph used, increasing the size of *X* could result in high degree nodes close to vertices in *P*, with the consequence discussed above.

**Figure S 1:**
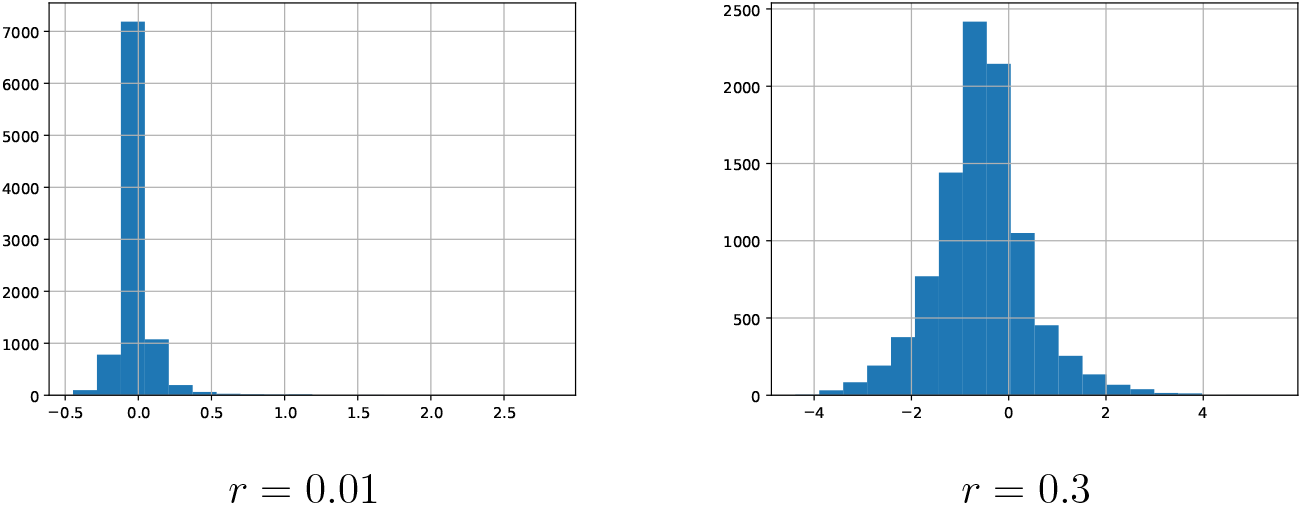
MINT: histogram of log scores for direction *X* ⇝ *P*.

**Figure S 2:**
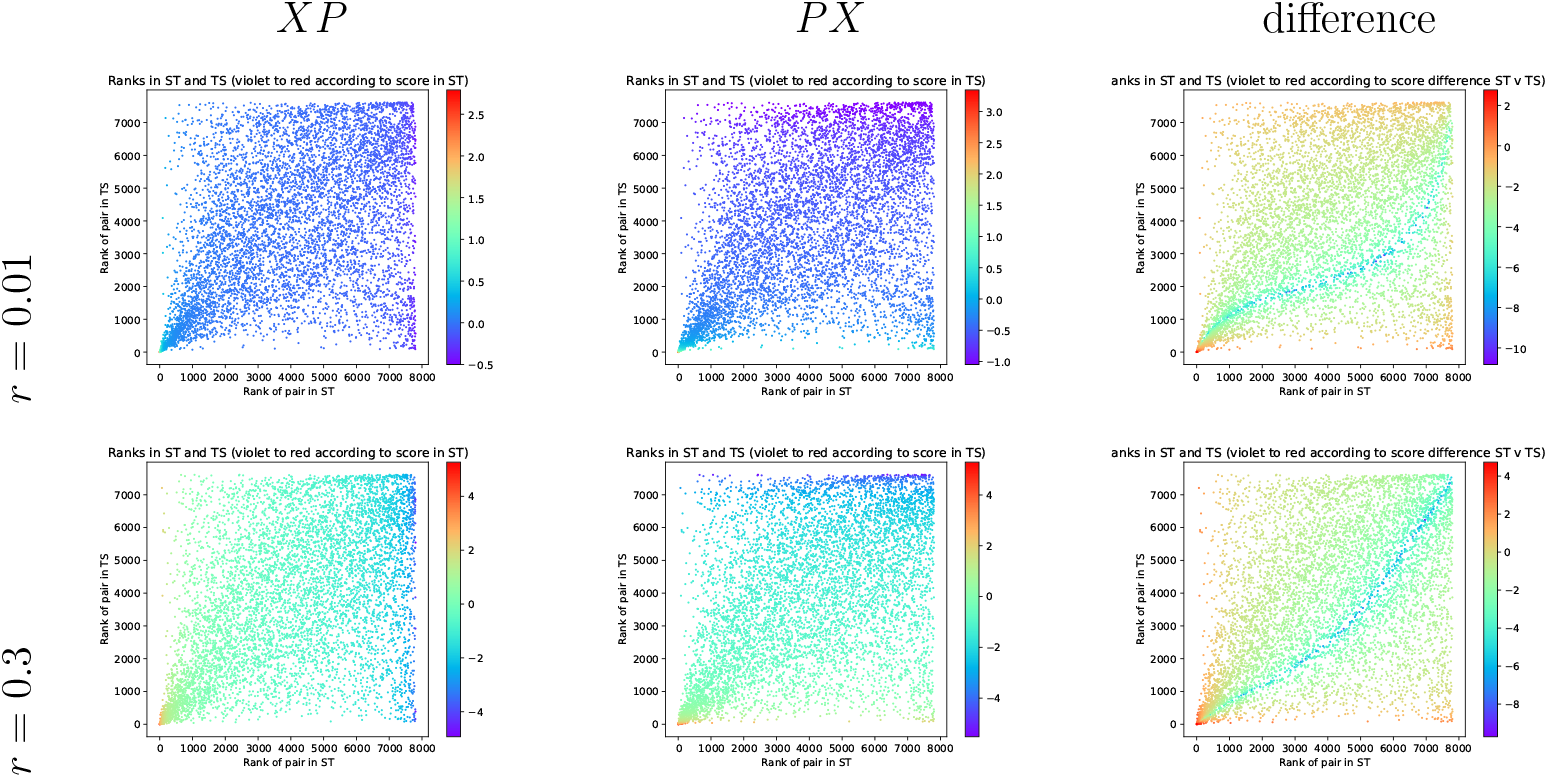
MINT: Scatter plot of log scores for the directions *X* ⇝ *P* and *P* ⇝ *X* Color coding of point is as follows: **(Left)** log *Q*^(*r*)^(*x,p*) **(Middle)** log *Q*^(*r*)^(*x,p*) **(Right)** log |*Q*^(*r*)^(*x,p*) — *Q*^(*r*)^ (*p,x*)|

### 7.2 Saturation indices and hits

**Figure S 3:**
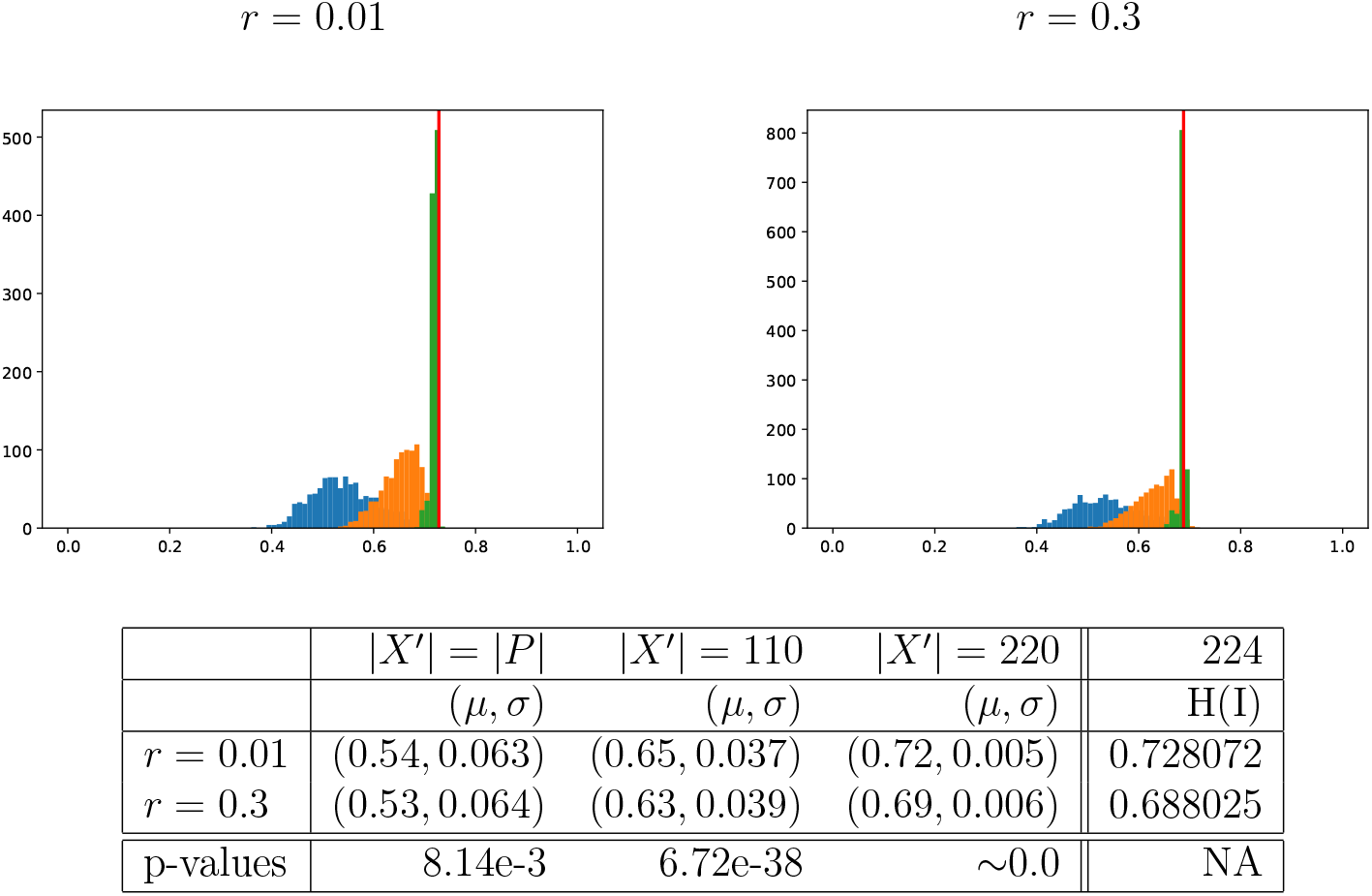
Symmetry score ratio *H*(*I*) (Eq. 7) at the instance level. Subsets *X*′ ⊂ *X* of size *s* ∈ {|*P*|, 110, 220} are used, *N_r_* = 1000 repeats for each size. **(Figures)** Distributions of the statistic *H*(*I*) for two restart rates *r* = 0.01 and *r* = 0.3: blue (|*X*′| = |*P*|) orange (|*X*′| = 110) green (|*X*′| = 220). Red lines represent the values of *H*(*X*). **(Table)** summaries *μ* and *σ* for *H*(*I*). The p-value reported is that of the Mann-Whitney U test, two-sided alternative. The last column of the table corresponds to the complete gene set *X*.

**Figure S 4:**
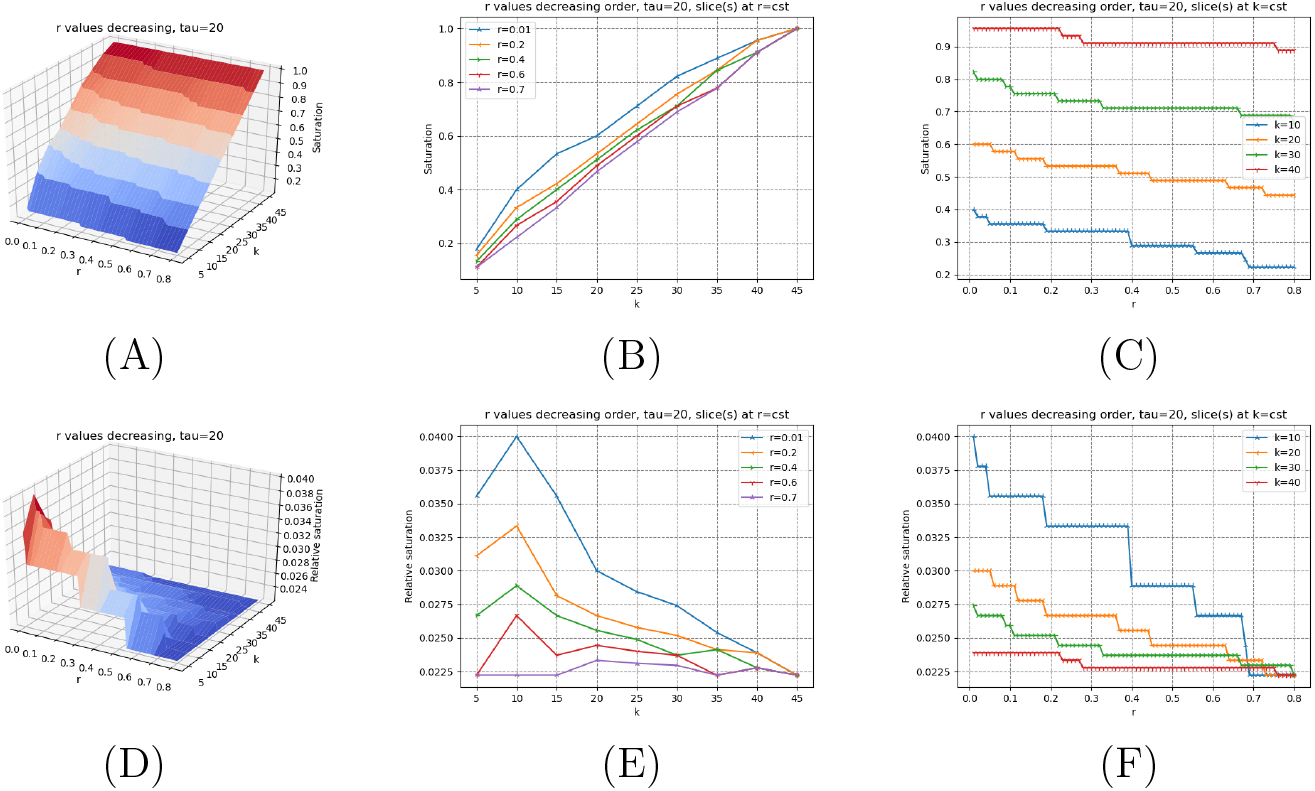
(Genetrank-renorm) Saturation plots (Def. 5) for *τ* = 20. Values of *r* processed in decreasing order. **(A, B, C)** Saturation index and slices at *r* = *cst* and *k* = *cst* (See Eq. 9) **(D, E, F)** Relative saturation index and slices at *r* = *cst* and *k* = *cst* (See Eq. 10)

**Figure S 5:**
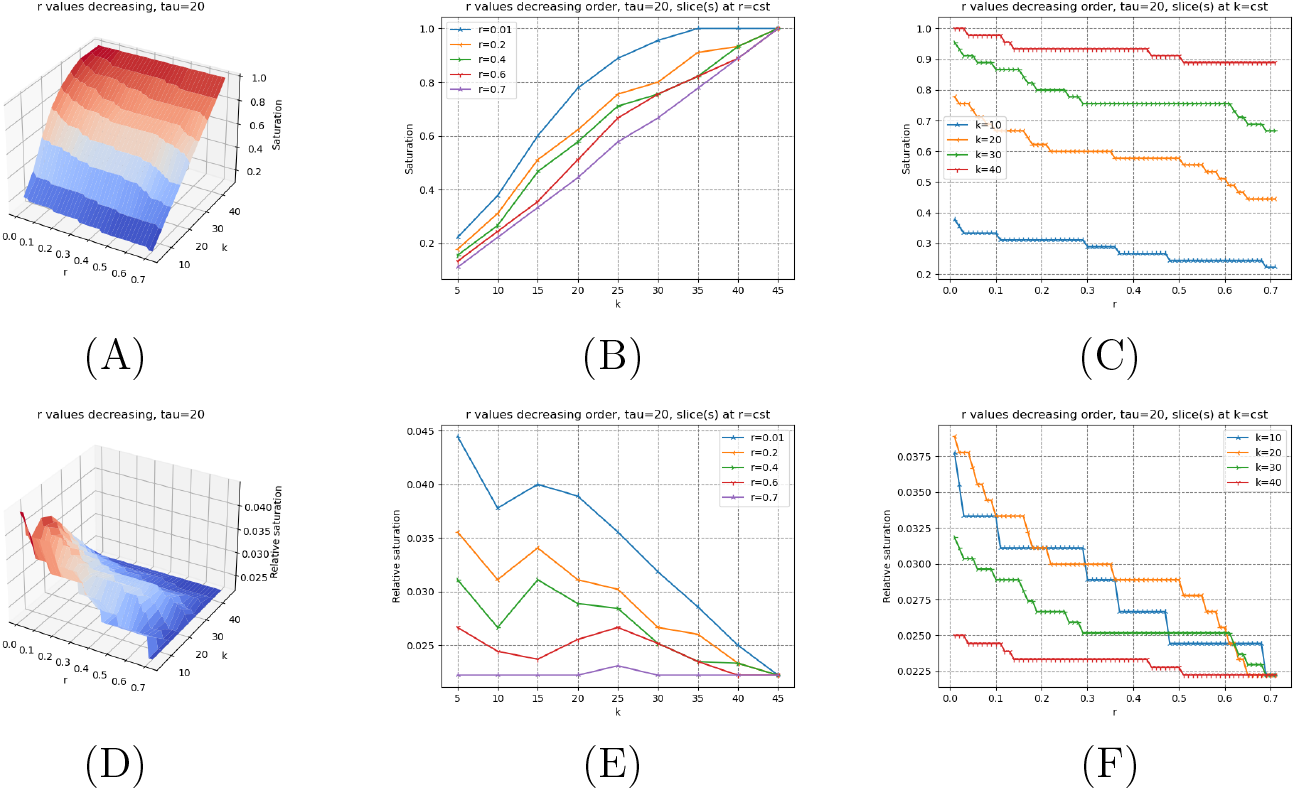
(Genetrank-AS) Saturation plots (Def. 5) for *τ* = 20. Values of *r* processed in decreasing order. **(A, B, C)** Saturation index and slices at *r* = *cst* and *k* = *cst* (See Eq. 9) **(D, E, F)** Relative saturation index and slices at *r* = *cst* and *k* = *cst* (See Eq. 10)

**Figure S 6:**
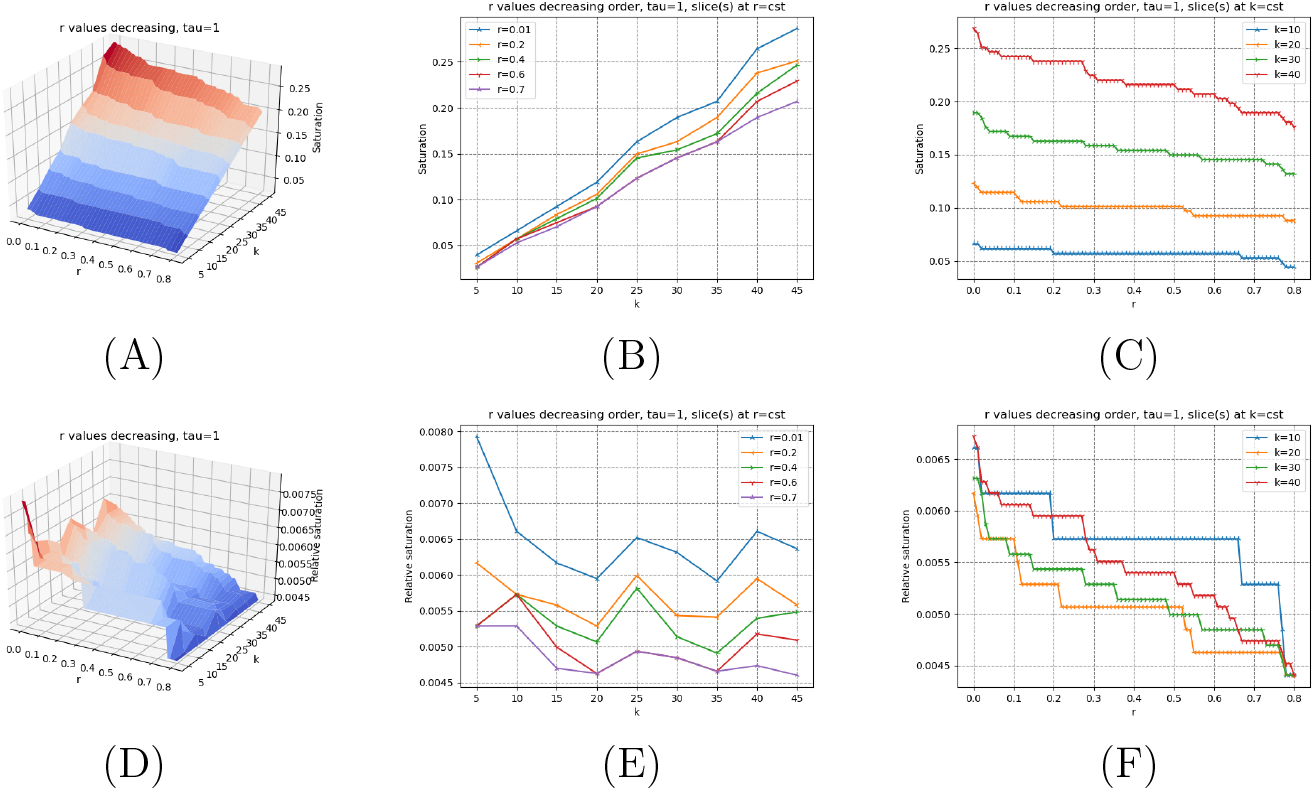
(Genetrank) Saturation plots (Def. 5) for *τ* = 1. Values of *r* processed in decreasing order. **(A, B, C)** Saturation index and slices at *r* = *cst* and *k* = *cst* (See Eq. 9) **(D, E, F)** Relative saturation index and slices at *r* = *cst* and *k* = *cst* (See Eq. 10)

**Figure S 7:**
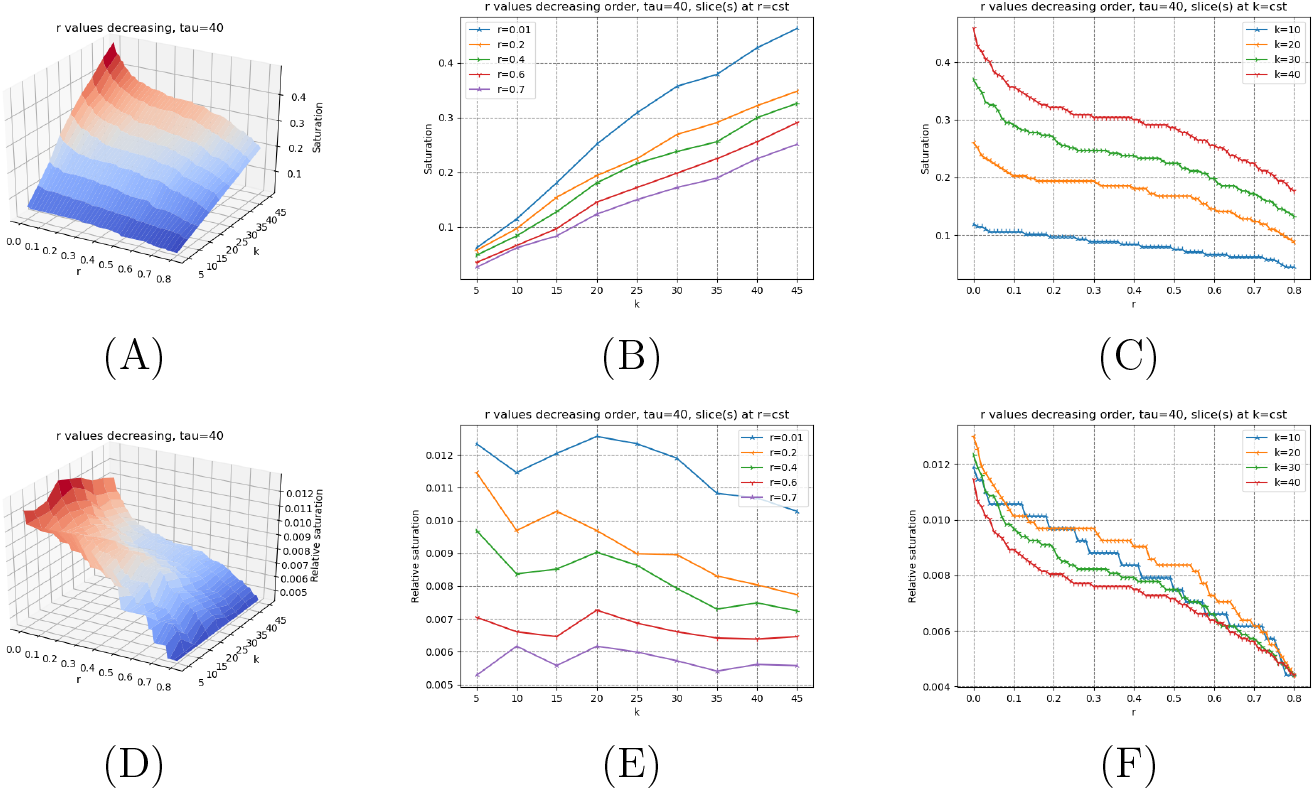
(Genetrank) Saturation plots (Def. 5) for *τ* = 40. Values of *r* processed in decreasing order. **(A, B, C)** Saturation index and slices at *r* = *cst* and *k* = *cst* (See Eq. 9) **(D, E, F)** Relative saturation index and slices at *r* = *cst* and *k* = *cst* (See Eq. 10)

### 7.3 Hits

**Figure S 8:**
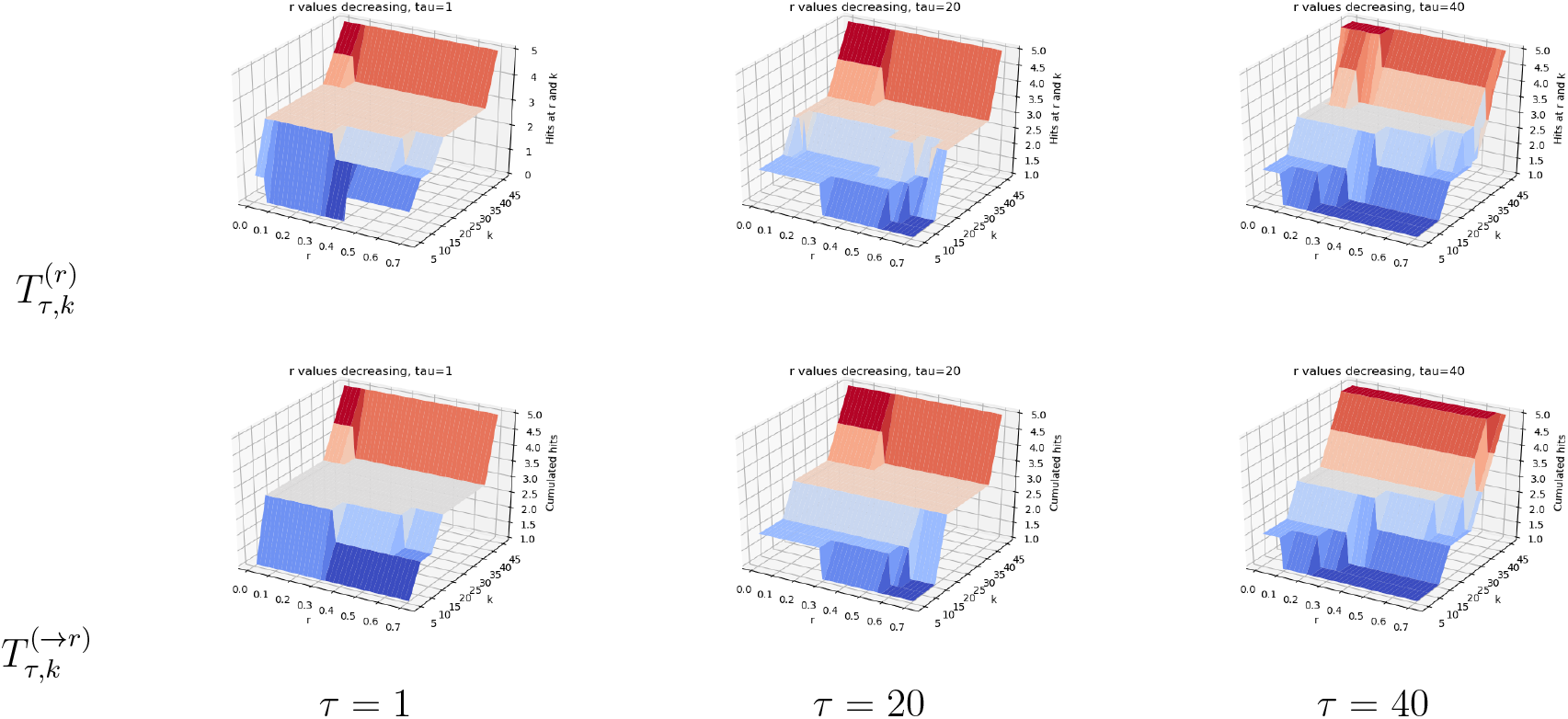
(Genetrank-renorm) Hits (Def. 6) for the list of reference genes O15304 (SIVA1), P78537 (BLOC1S1), P0CW18 (PRSS56), P53007 (SLC25A1), Q9Y2X8 (UBE2D4), DNM1L(O00429).

**Figure S 9:**
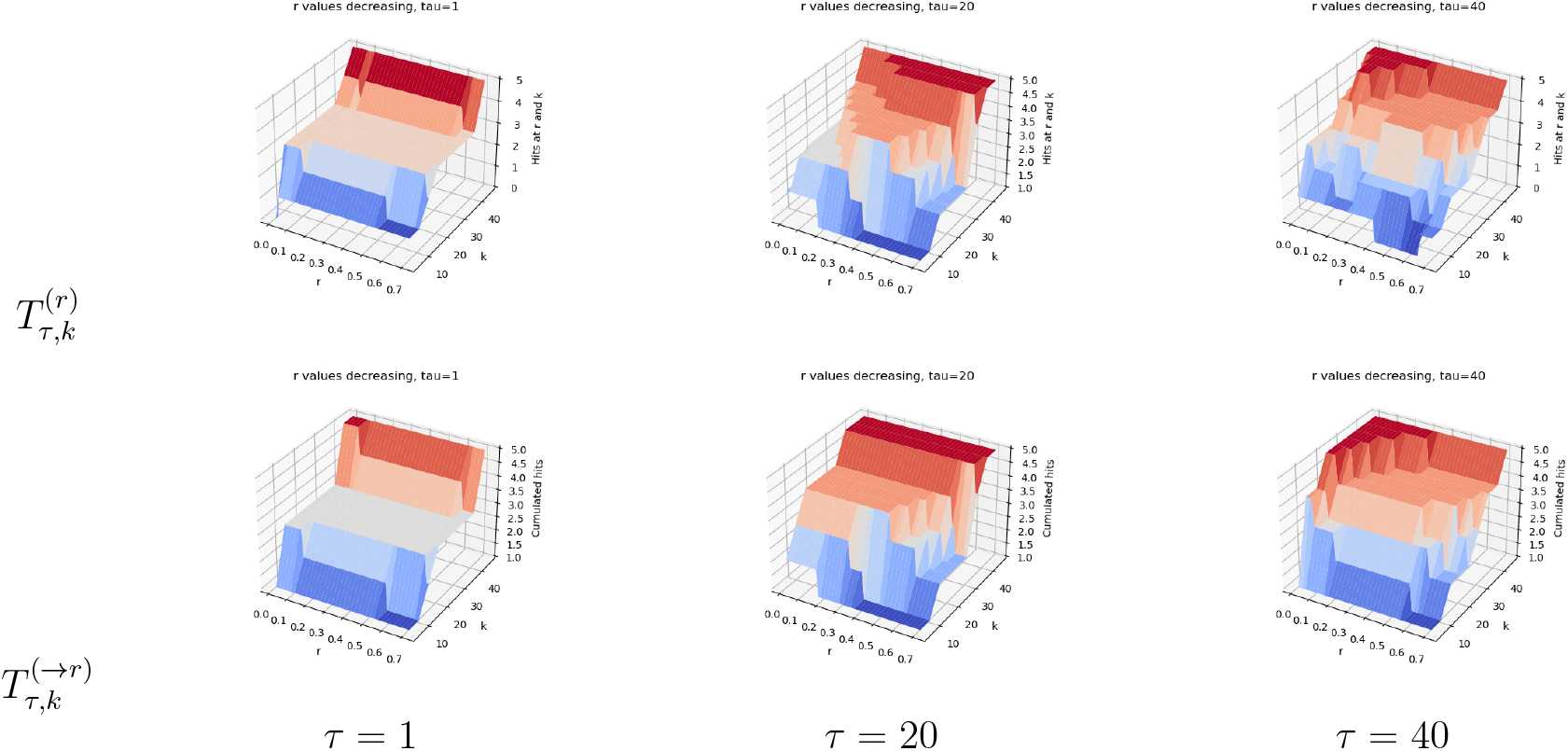
(Genetrank-AS) Hits (Def. 6) for the list of reference genes O15304 (SIVA1), P78537 (BLOC1S1), P0CW18 (PRSS56), P53007 (SLC25A1), Q9Y2X8 (UBE2D4), DNM1L(O00429).

### 7.4 Deferentially expressed genes

**Table S 1:**
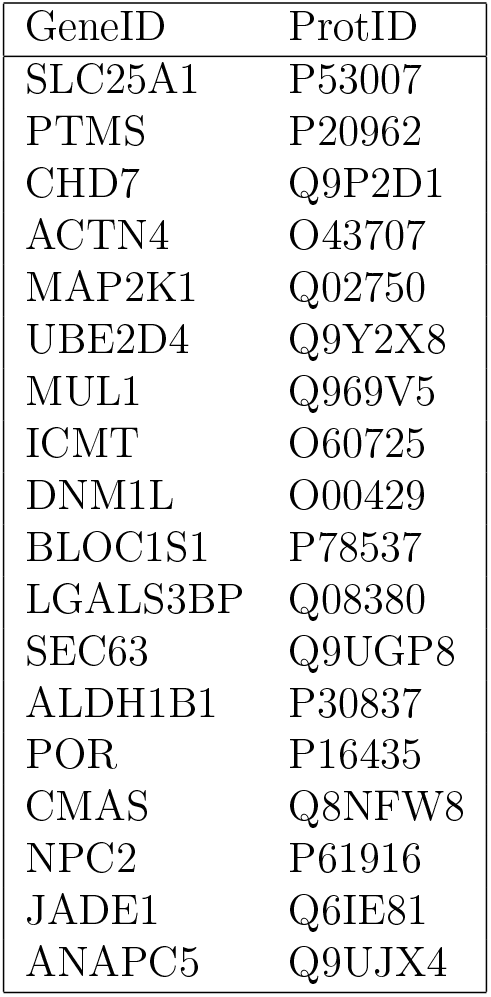
Set of 18 genes obtained by intersecting the list of 65 genes yielded by edgeR, with this list yielded by Genetrank (*k* = 50, range of values of *r*: 0..0.8, *τ* = 41).

7.4 Differentially expressed genes
| GeneID | ProtID |
| --- | --- |
| SLC25A1 | P53007 |
| PTMS | P20962 |
| CHD7 | Q9P2D1 |
| ACTN4 | O43707 |
| MAP2K1 | Q02750 |
| UBE2D4 | Q9Y2X8 |
| MUL1 | Q969V5 |
| ICMT | O60725 |
| DNM1L | O00429 |
| BLOC1S1 | P78537 |
| LGALS3BP | Q08380 |
| SEC63 | Q9UGP8 |
| ALDH1B1 | P30837 |
| POR | P16435 |
| CMAS | Q8NFW8 |
| NPC2 | P61916 |
| JADE1 | Q6IE81 |
| ANAPC5 | Q9UJX4 |

## Footnotes

1 Because throughout this paper we make use of a single PPIN (MINT) and the same set P, we will use the shorthand (*X*) for the instance.

## Notes

https://sbl.inria.fr/doc/Genetrank-user-manual.html

### Competing Interest Statement

The authors have declared no competing interest.

### Summary of Updates

The revision provides a better positioning with respect to existing work, stressing the difficulties of interaction networks targeted by our random walks with absorbing states.

